# Early-life vancomycin and meropenem treatment exert distinct effects on microbiota-gut-brain axis signalling in zebrafish larvae

**DOI:** 10.64898/2026.10.08.757590

**Authors:** Isabella Faimann, Annesha Ghosh, Eva Tatzl, Nastasia Stojanovic, Romana Penker, Giovanny Rodriguez-Blanco, Florian Reichmann

## Abstract

The microbiota-gut-brain axis is a bidirectional communication pathway between the gastrointestinal tract, gut microbiota and the central nervous system, that influences host neurobiology, metabolism and behaviour. Although germ-free and antibiotic-treated animals demonstrate a role for microbial signalling in mood and behaviour, mechanisms linking microbiome disruption to altered neural signalling during early development remain incompletely understood. Here, we investigated the impact of antibiotic-induced perturbations of the gut microbiota on behaviour, neurotransmitter levels as well as central and peripheral metabolism in 5 dpf zebrafish larvae, an established neurodevelopmental model. Larvae were exposed to the broad-spectrum antibiotic meropenem, the Gram-positive-directed antibiotic vancomycin, or their combination, and assessed gut microbiota composition, behaviour, neurotransmitter levels and metabolite profiles. All treatments altered microbial community composition, although the extent differed markedly, with pronounced restructuring after meropenem and combination treatment and more subtle changes after vancomycin. Behavioural analysis using the light-dark transition assay revealed reduced centre locomotion during the dark phase in all treated groups, without a general reduction in swimming activity, consistent with anxiety-like behaviour. These behavioural changes were accompanied by reduced expression of serotonergic genes and widespread central and peripheral metabolic disruptions in all antibiotic-treated groups. Treatment- and tissue-specific effects included suppression of the arginine-polyamine pathway in the brain following vancomycin treatment and increased peripheral glutamate, GABA and L-DOPA levels after meropenem exposure. Together, these findings demonstrate that early-life microbiome perturbation reshapes behaviour, neurotransmitter levels, and host metabolism in zebrafish larvae. In addition, we identified candidate metabolic pathways through which the microbiome may influence neurodevelopment.

## Introduction

The gut microbiome plays a crucial role in maintaining host health by regulating digestion, metabolic processes, immune responses, and communication between the gut and the brain (Cryan et al., 2019). It consists of a dynamic consortium of microorganisms that influence various physiological functions through interactions with host cells and the production of signalling molecules (Basnet et al., 2019; Gaulke et al., 2020; Li et al., 2022; Zhong et al., 2022; Sree Kumar et al., 2023). Understanding the complex interactions between the microbiome and host physiology has become a key area of biomedical research, with model organisms providing valuable insights into these relationships.

Zebrafish (*Danio rerio*) are a well-established vertebrate model for gut microbiome research due to several advantages of this animal model (Xia et al., 2022; Zhong et al., 2022). For example, zebrafish are highly fertile and display rapid development and optical transparency in larval stages, which permits high throughput screenings at early life stages (Stagaman et al., 2020). In addition, their lack of a functional adaptive immune system in early stages permits the investigation of fundamental microbial interactions without immune interference (Basnet et al., 2019; Levraud et al., 2022). Zebrafish share conserved physiological and genetic pathways with mammals, making them suitable for translational research, drug screenings, and studies on host-microbiome interactions (Gaulke et al., 2020; Rosa et al., 2022). Specifically, recent studies using zebrafish have begun to investigate how alterations in the gut microbiome influence host neurodevelopment and behaviour, highlighting the utility of this model for gut-brain axis research (Kiran et al., 2024). For example, dietary administration of the probiotic *Lactobacillus rhamnosus* IMC 501 to adult zebrafish modulates shoaling patterns and alters the expression of neurotrophic genes and genes of the serotonergic neurotransmitter system, indicating microbiota-driven changes in neurochemistry and behaviour (Borrelli et al., 2016). Moreover, supplementation with *Lactobacillus plantarum* reduced anxiety-like behaviour in adult zebrafish, altered GABAergic and serotonergic signalling in the brain and protected against stress-induced dysbiosis (Davis, Doerr, et al., 2016). Conversely, disruption of the gut microbiome through antibiotic exposure or rearing in germ-free conditions has been shown to alter locomotor activity in zebrafish larvae, demonstrating that microbial colonization during early development is essential for normal neurobehavioural outcomes (Phelps et al., 2017). Germ-free larvae additionally showed altered anxiety-related behaviour and failed to mount the characteristic response to an acute stressor, indicating that the absence of microbial colonization affects stress reactivity as well as baseline activity (Davis, Bryda, et al., 2016) . These findings collectively support the concept that both positive (probiotic) and negative (antibiotic/germ-free) perturbations of the gut microbiome can impact behavioural phenotypes in zebrafish, through changes in neurochemical pathways relevant to the gut-brain axis.

Despite this growing body of work, the mechanisms by which specific microbial perturbations influence gut-brain signalling in zebrafish, particularly at early developmental stages, remain poorly understood. Antibiotic-induced dysbiosis has been characterised in zebrafish, but almost exclusively using compounds relevant to aquaculture and environmental contamination such as tetracyclines, sulfonamides, fluoroquinolones and phenicols, typically at sub-therapeutic, environmentally relevant concentrations (Almeida et al., 2019; Kayani et al., 2021; Pindling et al., 2022; Yu et al., 2023). For example, subclinical streptomycin reduces microbial diversity in larvae (Pindling et al., 2022), and early-life oxytetracycline exposure alters gut community composition alongside larval hyperactivity and reduced midbrain neuron numbers (Yu et al., 2023). What has not been addressed though is how clinically used antibiotics with contrasting antimicrobial spectra reshape the larval microbiome, and whether the resulting communities influence gut-brain signalling to alter behaviour and neurochemistry.

To address this gap, the present study investigated how the widely used antibiotics vancomycin and meropenem as well as their combination influence microbiome composition and gut-brain axis signalling in zebrafish larvae. The two agents were selected because they show large differences in their antimicrobial spectrum. Vancomycin is primarily active against aerobic and anaerobic Gram-positive bacteria, while meropenem is active against a wide range of Gram-positive, Gram-negative, and anaerobic bacteria (Levine, 2006; Zhanel et al., 2007). Community composition was characterised by 16S rRNA gene sequencing and complemented by metabolomic profiling, brain and body neurotransmitter quantification and behavioural phenotyping, allowing compositional changes to be related directly to neurochemical and behavioural outcomes. We hypothesized that antibiotics with contrasting antimicrobial spectra differentially affect microbiome composition, early-life neurobehavioural development and signalling pathways including major neurotransmitters and metabolic responses.

## Results

### Vancomycin and meropenem exert distinct effects on gut microbial diversity, composition, and predicted function in zebrafish larvae

As a first step, we investigated how the chosen antibiotics affect the larval gut microbiome. Microbial community profiling using 16S rRNA sequencing showed that both meropenem and vancomycin as well as their combination markedly altered microbial diversity and composition (Fig. 1). Specifically, the Chao1 index (Fig. 1A), Shannon diversity (Fig. 1B), and Simpson index (Fig. 1C) were all significantly elevated in the meropenem and the meropenem/vancomycin combination group relative to control (*p* < 0.001), suggesting enhanced alpha diversity after antibiotic administration. In contrast, vancomycin did not change alpha diversity compared to the control group (Fig. 1A-C). Beta-diversity analysis also demonstrated clear separations between the treatment groups (Fig. 1D-F). Principal coordinates analysis (PCoA) based on Bray-Curtis dissimilarity and weighted UniFrac analysis revealed that meropenem- and meropenem/vancomycin-treated samples form distinct clusters from the control and vancomycin group along PC1 of the PCoA plot (Fig. 1D, E), suggesting strong changes in relative abundance of taxa due to meropenem. Unweighted UniFrac analysis likewise separated meropenem and meropenem/vancomycin samples from controls, but in addition revealed a partial separation of the vancomycin group (Fig. 1F), indicating treatment-specific differences in community membership rather than in the abundance of dominant taxa. To better characterize the effect of the chosen antibiotics, we next analysed differences in taxonomic composition. At the phylum level, microbiome composition in the control group was strongly dominated by *Proteobacteria* (Fig. 1G). Both vancomycin and meropenem administration led to an average reduction in relative *Proteobacteria* abundance, although the effect was much more pronounced in the meropenem and the meropenem/vancomycin combination group. We also found that as a consequence of this the relative abundance of various other phyla increased in the meropenem and the combination group including *Bacteroidota*, *Firmicutes* and *Actinobacteriota* (Fig. 1G). At the genus level, we found that the microbiota of control larvae was dominated by *Pseudomonas, Achromobacter* and *Aeromonas* (Fig. 1H). Vancomycin treatment led to a very strong reduction of *Aeromonas* and a rise in Pseudomonas, while meropenem and the combination treatment reduced all dominant taxa found in control larvae and increased the abundance of numerous other bacterial genera (Fig. 1H).

**Figure 1:**
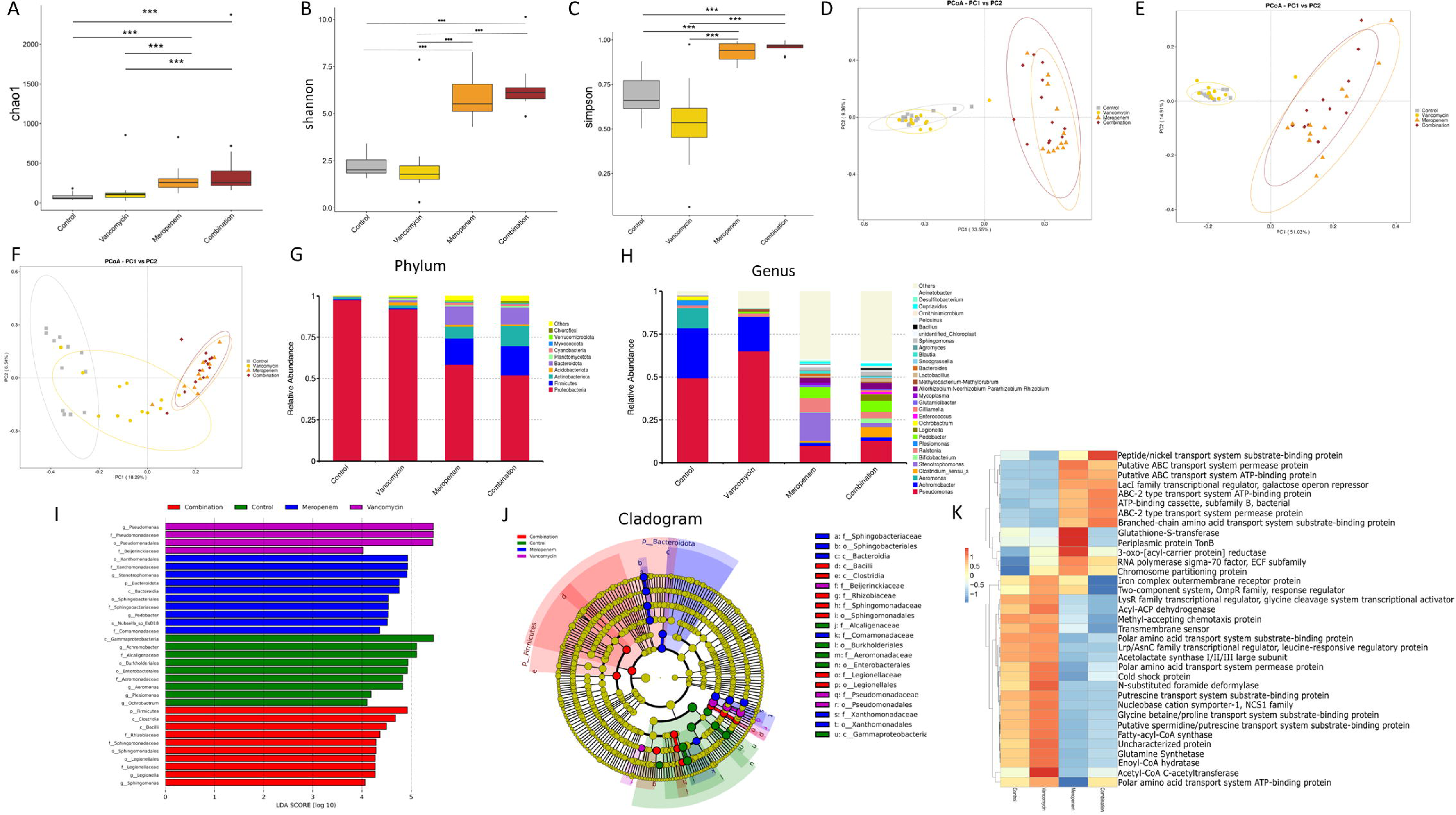
Antibiotic treatment alters microbial diversity, composition, and predicted functional gene content in zebrafish larvae. Microbiome profiling of zebrafish larvae following exposure to vancomycin, meropenem, or a combination of both antibiotics in comparison to unexposed larvae (control). (**A-C**) Alpha-diversity indices (Chao1, Shannon, and Simpson) showing significant increases in richness and diversity in the meropenem and combination treated groups. (**D-F**) Principal coordinates analysis (PCoA) based on (**D**) Bray-Curtis dissimilarity, (**E**) weighted UniFrac and (**F**) unweighted UniFrac showing separation of meropenem-and combination-treated samples from control and vancomycin samples along PC1. Ellipses represent 95% confidence intervals. (**G,H**) Relative abundance of **(G)** the 10 most abundant bacterial phyla and **(H)** the 30 most abundant bacterial genera in the study groups showing shifts in dominant taxa following antibiotic exposure. The less abundant remaining taxa are displayed as “Others”. (**I**) LEfSe cladogram depicting differentially enriched bacterial taxa among treatment groups (LDA score > 4.0). (**J**) Bar plot of LDA scores of significantly different taxa among treatment groups (LDA score > 4). (**K**) Heatmap of significant predicted KEGG Ortholog (KO) abundances inferred using PICRUSt2. Colours represent row-wise z-scores of group mean relative abundances and rows are ordered by hierarchical clustering. Each biological replicate represents a pool of six zebrafish larvae. Data in A-C are presented as boxplots (median, IQR, min-max). ***p<0.001, one-way ANOVA followed by Tukey post hoc test. Beta-diversity was assessed by PERMANOVA on Bray-Curtis, weighted and unweighted UniFrac distances. LEfSe analysis consisted of Kruskal-Wallis across classes, pairwise Wilcoxon rank-sum test and LDA threshold 4. For PICRUSt2 differences in predicted KO abundances between groups were assessed by Kruskal-Wallis test with Benjamini-Hochberg correction (q < 0.05). p < 0.05 was considered statistically significant. n = 10 replicates per treatment group.

To assess statistically significant differentially abundant taxa between the experimental groups, we used LEfSe (Linear discriminant analysis Effect Size) analysis (Fig. 1I,J). We found that control larvae show a significant enrichment of multiple members of the *Proteobacteria* phylum, including *Gammaproteobacteria*, *Aeromonadaceae*, *Enterobacterales*, Achromobacter, Alcaligenaceae and *Burkholderiales*, consistent with the *Proteobacteria*-dominated microbiome typical of early-life zebrafish (Li et al., 2022; Garibay-Valdez et al., 2024). Vancomycin treatment was also characterized by a significant enrichment of *Proteobacteria*, particularly the Pseudomonas lineage (*Pseudomonadales*, *Pseudomonadaceae* and *Pseudomonas*) as well as *Beijerinckiaceae*, suggesting treatment-induced shifts within the *Proteobacteria* lineage. Meropenem treatment resulted in a distinct enrichment profile, with higher abundances of the *Bacteroidota* lineage (phylum to genus level, including Sphingobacteriales and Pedobacter), and of *Proteobacteria* lineages such as *Xanthomonadaceae* and *Comamonadaceae*. The Xanthomonadaceae signal was driven by Stenotrophomonas, an intrinsically carbapenem-resistant genus (Gil-Gil et al., 2020), which reached around 15% relative abundance in meropenem-treated larvae (Fig. 1H). In contrast to vancomycin, meropenem produced therefore both a strong selection for antibiotic-resistant taxa and broader phylogenetic turnover including taxa dominant in control larvae. The combination treatment group showed an enrichment profile largely overlapping with that of the meropenem group (Fig. 1I,J), including multiple clades within *Firmicutes* (*Bacilli* and *Clostridia*) and *Proteobacteria* (*Sphingomonas* lineage, *Legionella* lineage and *Rhizobiaceae*). Notably, the relative expansion of Firmicutes was of comparable magnitude in the meropenem-only group (Fig. 1G), indicating that it is primarily driven by the meropenem-induced loss of the dominant Gram-negative taxa rather than by the addition of vancomycin. To directly compare treatments, we additionally performed pairwise LEfSe analyses for all group combinations (Fig. S1). Comparison of vancomycin-treated larvae with controls confirmed the depletion of *Aeromonas*, *Aeromonadaceae*, *Plesiomonas* and *Enterobacterales* and the enrichment of the Pseudomonas lineage and Beijerinckiaceae (Fig. S1A), while no Gram-positive taxon was significantly enriched in controls. Comparisons of meropenem and of the combination treatment with controls yielded substantially larger numbers of differentially abundant taxa, including Firmicutes, Bacteroidota and Actinobacteriota lineages as well as Stenotrophomonas (Fig. S1B,C). Comparisons of the vancomycin group with the two meropenem-containing groups largely mirrored the corresponding control comparisons (Fig. S1D, E). Vancomycin-treated larvae showed higher relative abundance of the *Pseudomonas* lineage, *Achromobacter* and *Alcaligenaceae*, while *Firmicutes*, *Bacteroidota* and *Actinobacteriota* lineages, *Stenotrophomonas* and *Sphingomonadales* were enriched in the meropenem and combination groups. The *Legionella* lineage was enriched in the combination group relative to vancomycin (Fig. S1E) but did not differ between the vancomycin and meropenem groups (Fig. S1D). Direct comparison of the meropenem and combination groups revealed only seven differentially abundant taxa (Fig. S1F). *Stenotrophomonas*, *Xanthomonadaceae*, *Xanthomonadales* and *Comamonadaceae* showed higher relative abundance under meropenem alone, whereas the *Legionella* lineage was again enriched in the combination group, confirming the enrichment of the *Legionella* lineage in the combination group. Notably, *Firmicutes*, *Bacilli* and *Clostridia* did not differ significantly between these two groups, but were enriched in both relative to control and vancomycin (Fig. S1B,C,F).

PICRUSt2 functional prediction analysis revealed substantial differences in predicted microbial gene content across treatments (Fig. 1K). In line with the taxonomic composition differences, hierarchical clustering of KEGG orthologs (KOs) revealed a similar enrichment and depletion of pathway terms in the meropenem and meropenem/vancomycin combination group. In addition, we found that the control and the vancomycin group also showed a similar KO enrichment pattern, but one that differed significantly from the other two groups. Meropenem and meropenem/vancomycin combination exposure induced a shift in the predicted abundance of microbial genes related to transport systems with a predicted enrichment of ABC transporter genes, while other transporter systems had a lower predicted abundance. This includes genes related to polyamine transport (spermidine and putrescine transport system substrate-binding protein) and genes related to amino acid and osmolyte transport (e.g., glycine betaine/proline transport, polar amino acid transport system substrate-binding protein). In addition, glutamine synthetase, an enzyme involved in glutamate metabolism, had also lower predicted abundance in the two meropenem groups. A unique feature of the meropenem group is the predicted higher abundance of glutathione-S-transferase, periplasmic protein TonB, and 3-oxoacyl-[acyl-carrier protein] reductase compared to all other treatment groups and a unique feature of the combination group is the predicted depletion of iron complex outer membrane receptor and the OmpR-family two-component regulator. Notably, TonB, the energy transducer for outer membrane iron transport, showed higher predicted abundance under meropenem, whereas the iron complex outer membrane receptor, a Ton-B dependent receptor (Noinaj et al., 2010), showed lower predicted abundance in the combination group, suggesting differences in predicted iron uptake capacity. As mentioned above, the vancomycin group was similar to the control group, but the predicted abundance of enzymes involved in fatty acid β-oxidation including Acetyl-CoA C-acetyltransferase, fatty-acyl-CoA synthase, and enoyl-CoA hydratase was higher in this group compared to control conditions. Taken together, these findings suggest that antibiotic-induced community restructuring was accompanied by antibiotic-specific shifts in the predicted functional gene repertoire of the community. We found that meropenem exposure profoundly restructures the larval zebrafish microbiota, whereas vancomycin produces a more subtle shift. Both treatments nonetheless yielded distinct, treatment-specific microbial and predicted functional profiles.

### Vancomycin and meropenem exert treatment-specific effects on locomotion and anxiety in the light-dark transition assay

To study how the chosen antibiotics affect gut-brain axis signalling in zebrafish larvae, we next investigated larval behaviour after antibiotic administration. For this, we performed the light-dark transition test, an assay to evaluate larval locomotion and anxiety (Hillman et al. 2024). As shown in Fig. 2A, larvae from all groups displayed higher locomotor activity during dark phases compared to light phases (RM-ANOVA; *p* < 0.0001). However, detailed analysis per minute revealed that larvae in the meropenem/vancomycin combination group exhibited higher locomotor activity during the initial light phase compared to the controls and vancomycin group (Fig. 2A; *p* < 0.05) suggesting impaired habituation to the test conditions. When analysing the average distance moved in the arena during light and dark phases, we found again a rise in locomotion during the dark phase independent of antibiotic treatment (Fig. 2B; *p* < 0.0001). In the dark-flash analysis (locomotion directly after lights went off), all groups showed a rapid increase in locomotion immediately following dark onset (Fig. 2C; *p* < 0.0001), suggesting that the antibiotic treatments did not impair general locomotion or light perception. Analysis of locomotor activity in the inner zone of the test arena, a readout of anxiety-like behaviour, revealed a different pattern from that seen for total distance. Whereas all groups increased overall locomotion in the dark (Fig. 2B), only control and meropenem larvae also moved significantly more in the inner zone during dark phases than during light phases (Fig. 2D; *p* < 0.0001 and *p* = 0.039, respectively), indicating a dark-induced shift towards centre exploration consistent with reduced anxiety-like behaviour. Interestingly, this shift was absent in the vancomycin and the meropenem/vancomycin combination group (Fig. 2D; light condition X treatment interaction: *p* = 0.0248), which increased overall locomotion in the dark without a corresponding increase in the inner zone. Moreover, inner-zone locomotion during the dark phase was significantly lower in all three antibiotic-treated groups than in controls (Fig. 2D), indicating that antibiotic treatment attenuated the dark-associated centre exploration seen in untreated larvae, consistent with a treatment-associated increase in anxiety-like behaviour.

**Figure 2:**
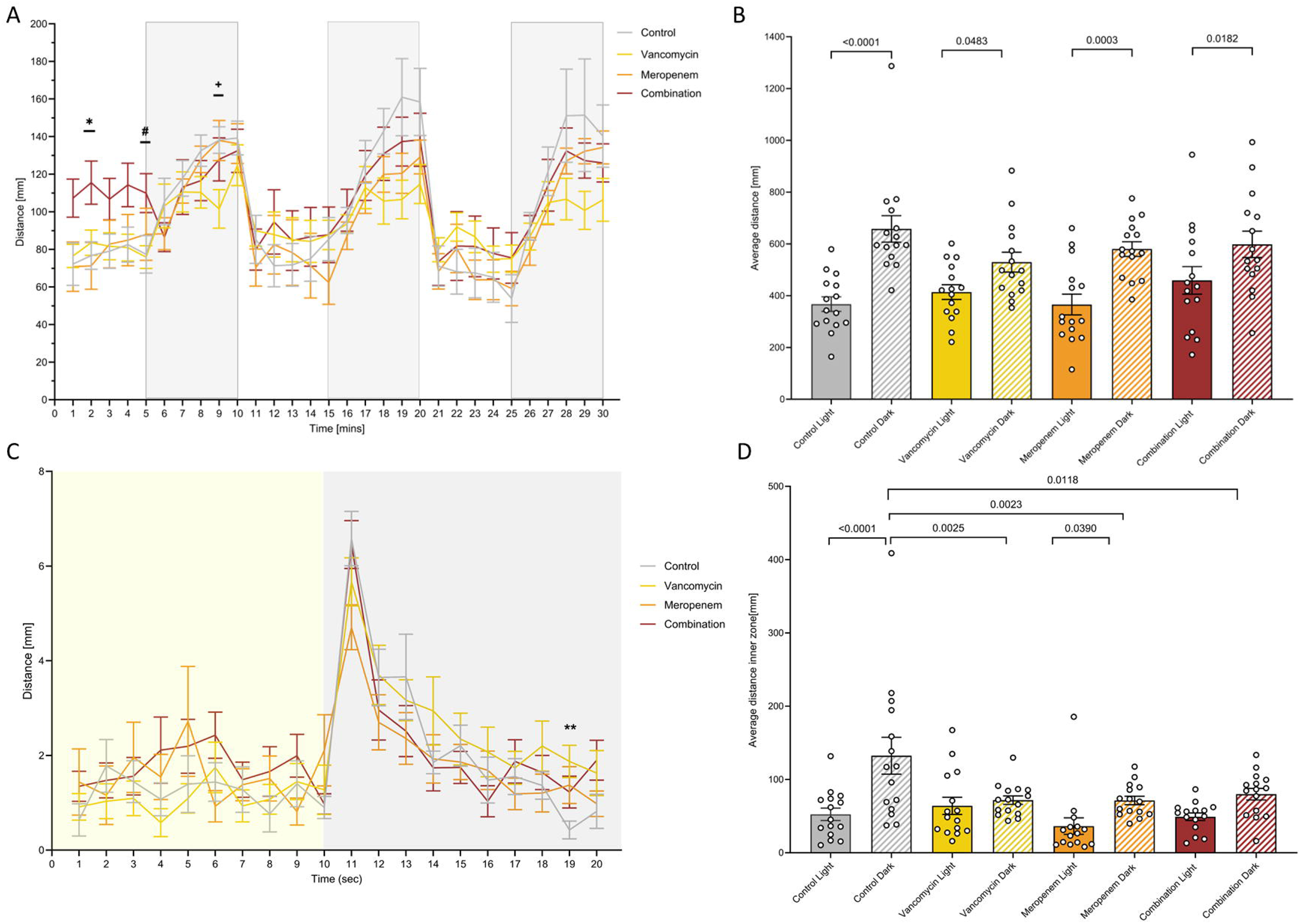
Antibiotic treatment alters larval locomotor and anxiety-like behaviour in the light-dark transition assay. (**A**) Per minute analysis of the total distance travelled in the test arena during the light-dark transition assay. Larvae were exposed to alternating light (white background) and dark (visible light off; grey background) phases. RM-ANOVA (time effect: F_(6.427, 359.9)_ = 20.54, p < 0.0001) followed by Tukey’s post-hoc test. *p < 0.05 control vs. meropenem/vancomycin combination; ^#^p < 0.05 vancomycin vs. meropenem/vancomycin combination; ^+^p < 0.05 control vs. vancomycin. Data are presented as mean ± SEM (n = 15 per group). (**B**) Average distance travelled in the test arena during light and dark phases. Two-way ANOVA (light condition main effect: F_(1, 112)_ = 42.84, p < 0.0001) followed by Tukey’s post-hoc test. Individual data points are overlaid on bar graphs showing mean ± SEM. n = 15 per group. (**C**) Distance travelled per second before and after lights go off for the first time (dark flash). The yellow background indicates the light phase and the grey background the dark phase. RM-ANOVA (time effect: F_(11.19, 626.9)_ = 20.92, p < 0.0001) followed by Tukey’s post-hoc test. **p < 0.01 control vs. vancomycin. Data are presented as mean ± SEM (n = 15 per group). (**D**) Average distance travelled in the inner zone of the test arena during light and dark phases. Two-way ANOVA (light condition × treatment interaction: F_(3, 112)_ = 3.241, p = 0.0248; light condition main effect: F_(1, 112)_ = 20.98, p < 0.0001; treatment main effect: F_(3, 112)_ = 3.789, p = 0.0124) followed by Tukey’s post-hoc test. Individual data points are overlaid on bar graphs showing mean ± SEM. n = 15 per group.

### Antibiotic exposure alters the serotonergic system in the larval zebrafish brain

Given that the observed behavioural alterations after antibiotic treatment are indicative of anxiety, we next investigated whether these changes in behaviour are accompanied by changes in the brain serotonergic system, an important regulator of anxiety across species (Millan, 2003; Maximino et al., 2013). Therefore, we next measured serotonin levels in larval zebrafish heads using a targeted metabolomics approach. Central 5-HT levels did not differ significantly from controls in any treatment group, although meropenem- and combination-treated larvae differed from one another, with higher levels in the meropenem group (Fig. 3A). In contrast, other central neurotransmitter levels were unaltered after treatment with meropenem, vancomycin or their combination (Table S1). Gene expression analysis revealed that the serotonin-synthesizing enzyme *tryptophan hydroxylase 1a* (*tph1a*) was significantly downregulated in all three antibiotic-treated groups relative to controls, suggesting that serotonin synthesis capacity is reduced, irrespective of the antibiotic used (Fig. 3B). By contrast, *tryptophan hydroxylase 2* (*tph2*), the isoform predominantly expressed in raphe serotonergic neurons, was not significantly altered (Fig. 3C). The expression of the serotonin transporter *solute carrier family 6 member 4a* (*slc6a4a*) was reduced in all treated groups, reaching statistical significance only in meropenem-treated larvae (Fig. 3D), whereas its paralogue *solute carrier family 6 member 4b* (*slc6a4b*) was unaffected by the antibiotic treatments (Fig. 3E). The expression of *monoamine oxidase* (*mao*), a key enzyme involved in serotonin degradation, was likewise not significantly altered (Fig. 3F). Taken together, these findings indicate that antibiotic exposure affected the serotonergic system, with a consistent transcriptional reduction in *tph1a* across all treatments, an additional meropenem-associated reduction in *slc6a4a*, as well as a difference in 5-HT levels between the meropenem and combination groups.

**Figure 3:**
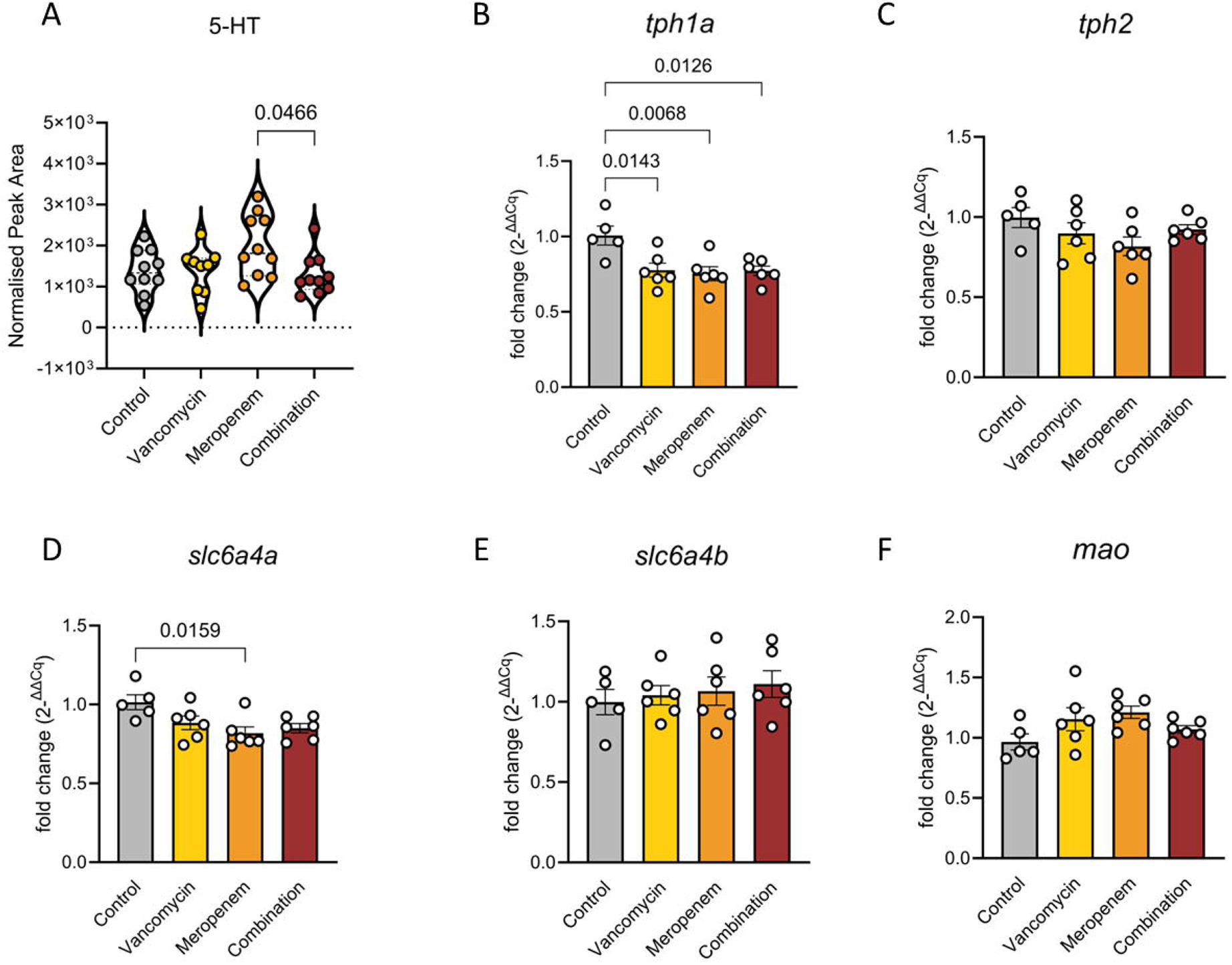
Antibiotic exposure alters the serotonergic system in the larval zebrafish brain. **(A)** Violin plot showing the normalised peak area of the 5-hydroxytryptamine (5-HT, serotonin) peak as measured by targeted metabolomics in head extracts of zebrafish larvae following treatment with vancomycin, meropenem, or vancomycin + meropenem (combination) compared to untreated larvae (control). Peak areas were normalised to sample protein content. Each dot represents a pool of five larval heads; violins indicate distribution density. One-way ANOVA followed by Tukey’s post hoc test. n = 8-10 replicates per treatment group. **(B-F)** Relative mRNA expression levels of serotonin-related genes in head samples of 5 days post-fertilization (dpf) zebrafish larvae following exposure to vancomycin, meropenem, or vancomycin + meropenem (combination) compared to untreated larvae (control). **(B)** tph1a (tryptophan hydroxylase 1a), **(C)** tph2 (tryptophan hydroxylase 2), **(D)** slc6a4a (solute carrier family 6 member 4a), **(E)** slc6a4b (solute carrier family 6 member 4b) and **(F)** mao (monoamine oxidase) expression. Expression levels were normalized to the reference genes actb1 and rpl13a and are presented as fold change (2^−ΔΔCq^) relative to controls. Data represent mean ± SEM, with each biological replicate consisting of a pool of five zebrafish larval heads. Statistical significance was determined using one-way ANOVA followed by Tukey post hoc test. n = 5-7 biological replicates per group.

### Metabolic effects of antibiotics on the brain of zebrafish larvae

Next, we investigated whether the chosen antibiotics affect neurotransmitter-independent pathways in larval heads using an extended metabolomic profiling method (Yuan et al., 2012). In total, we detected 85 metabolites using this method and found that 14 of these metabolites were significantly altered by one or more antibiotic treatments, seven of which also remained significant after false discovery rate correction across all comparisons (Table S2). Of particular note is the suppression of polyamine metabolism by the antibiotics, although these alterations did not remain significant in the more stringent corrected analysis. Specifically, within the arginine-polyamine axis, both arginine (Fig. 4A) and N-acetyl-L-ornithine, an intermediate in arginine and polyamine metabolism (Fig. 4B), were significantly reduced in vancomycin-treated animals compared to controls. Moreover, spermine, a downstream polyamine, was reduced in all 3 treatment groups compared to controls (Fig. 4C), whereas N-acetylputrescine, the only other polyamine-related metabolite detected, did not differ between groups (Table S2). Together with the vancomycin-specific reductions in arginine and N-acetyl-L-ornithine, this suggests that vancomycin treatment had a broader effect on this pathway than the other two treatments. In addition, metabolites involved in nucleotide biosynthesis and cellular energy metabolism were also significantly altered by the antibiotics, although again not after correction. 5-aminoimidazole-4-carboxamide ribonucleotide (AICAR), a purine synthesis intermediate and 5’ adenosine monophosphate-activated protein kinase (AMPK) activator, was significantly reduced in vancomycin-treated larvae relative to controls (Fig. 4D), while inosine, a purine catabolite, was significantly elevated in the combination treatment group compared to both controls and vancomycin-treated animals (Fig. 4E). We also found that biotin, a B-vitamin to which the gut microbiota can contribute, was significantly reduced in vancomycin-treated larvae relative to controls (Fig. 4F).

**Figure 4:**
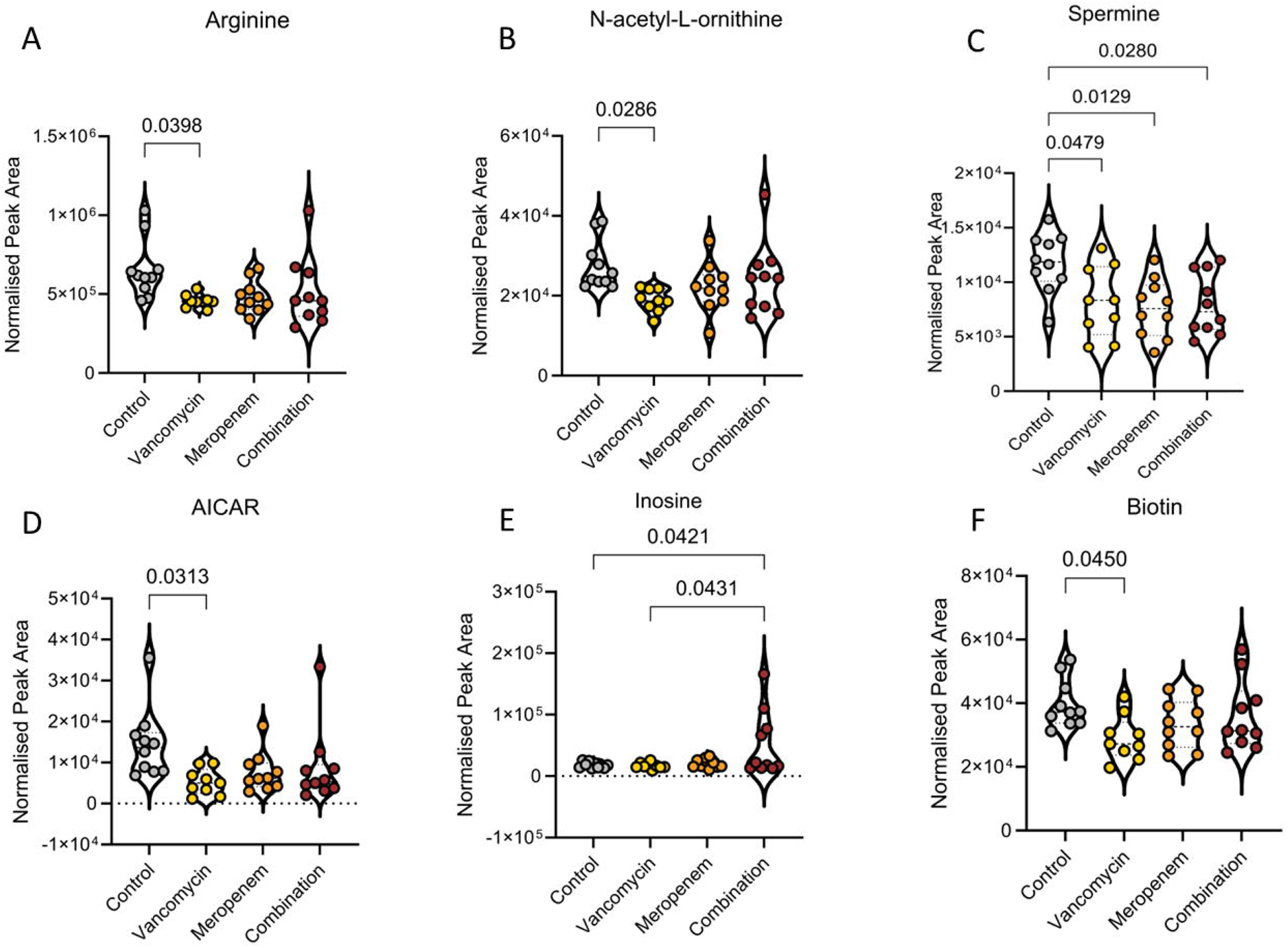
Treatment-specific metabolic alterations in larval zebrafish heads. **(A-F)** Violin plots showing normalised peak areas of metabolites significantly altered in head extracts of zebrafish larvae measured by metabolomics following treatment with vancomycin, meropenem, or vancomycin + meropenem (combination) compared to untreated larvae (control). (**A**) Arginine, (**B**) N-acetyl-L-ornithine, (**C**) Spermine, (**D**) 5-aminoimidazole-4-carboxamide ribonucleotide (AICAR), (**E**) Inosine and (**F**) Biotin. Each dot represents a pool of five larvae; violins indicate distribution density. One-way ANOVA followed by Tukey’s post hoc tests. n = 9-10 replicates per treatment group.

### Metabolites altered both in head and body compartments after antibiotic treatment

Given that antibiotic treatment had substantial effects on gut microbiota and the brain metabolome of larval zebrafish, we next performed metabolic measurements of larval body samples. We were able to detect the same metabolites as in zebrafish head samples and found that 32 of these metabolites were significantly altered between the experimental groups, with 22 also remaining significant in the more stringent analysis (Table S3). Eight metabolites were found to be significantly altered in both head and body compartments, seven of which remained significant after correction. Specifically, asparagine levels were elevated in both the head and body of meropenem-treated and meropenem/vancomycin combination-treated larvae compared to controls (Fig. 5A,B), suggesting a systemic disruption of amino acid homeostasis due to antibiotic treatment. Similar effects were observed for vancomycin, which also increased asparagine levels, but this reached significance only in body samples. In contrast, carbamoyl phosphate, a key intermediate in both the urea cycle and pyrimidine biosynthesis, was reduced in head samples after all antibiotic treatments (Fig. 5C) but increased in body samples only after meropenem treatment (Fig. 5D), suggesting a compartment-dependent disruption of nitrogen metabolism. Pipecolic acid, a lysine catabolite with neuromodulatory properties, was significantly increased in head samples after meropenem treatment only (Fig. 5E), but increased in body samples after both meropenem and meropenem/vancomycin combination treatment (Fig. 5F). Glycerophosphocholine, a membrane-derived choline metabolite, was profoundly decreased in head samples of all treatment groups (Fig. 5G), whereas in body samples it was increased after meropenem treatment but unaffected by vancomycin and combination treatment (Fig. 5H), suggesting antibiotic-induced disruption of phospholipid turnover that is particularly pronounced in head samples. In addition, hydroxyproline showed a significant increase in all antibiotic groups in both compartments investigated (Fig. 5I,J), but lysine, which was among the most robustly altered metabolites, showed the opposite pattern with markedly reduced levels in all treatment groups in both head and body (Fig. 5K,L), implicating the gut microbiome as a contributor to lysine availability in the larval zebrafish. Notably, lysine depletion was accompanied by increased pipecolic acid after meropenem and combination treatment (Fig. 5E,F), but not after vancomycin. NAD⁺ levels were decreased in head samples in the combination group only compared to all other treatment groups (Fig. 5M), but increased in body samples after meropenem treatment (Fig. 5N), indicating treatment- and compartment-specific effects on redox balance. Finally, phosphorylcholine was significantly reduced in head samples after vancomycin and meropenem treatment (Fig. 5O), while it was increased in body samples only after combination treatment (Fig. 5P). This was the only one of the eight metabolites that did not remain significant after correction.

**Figure 5:**
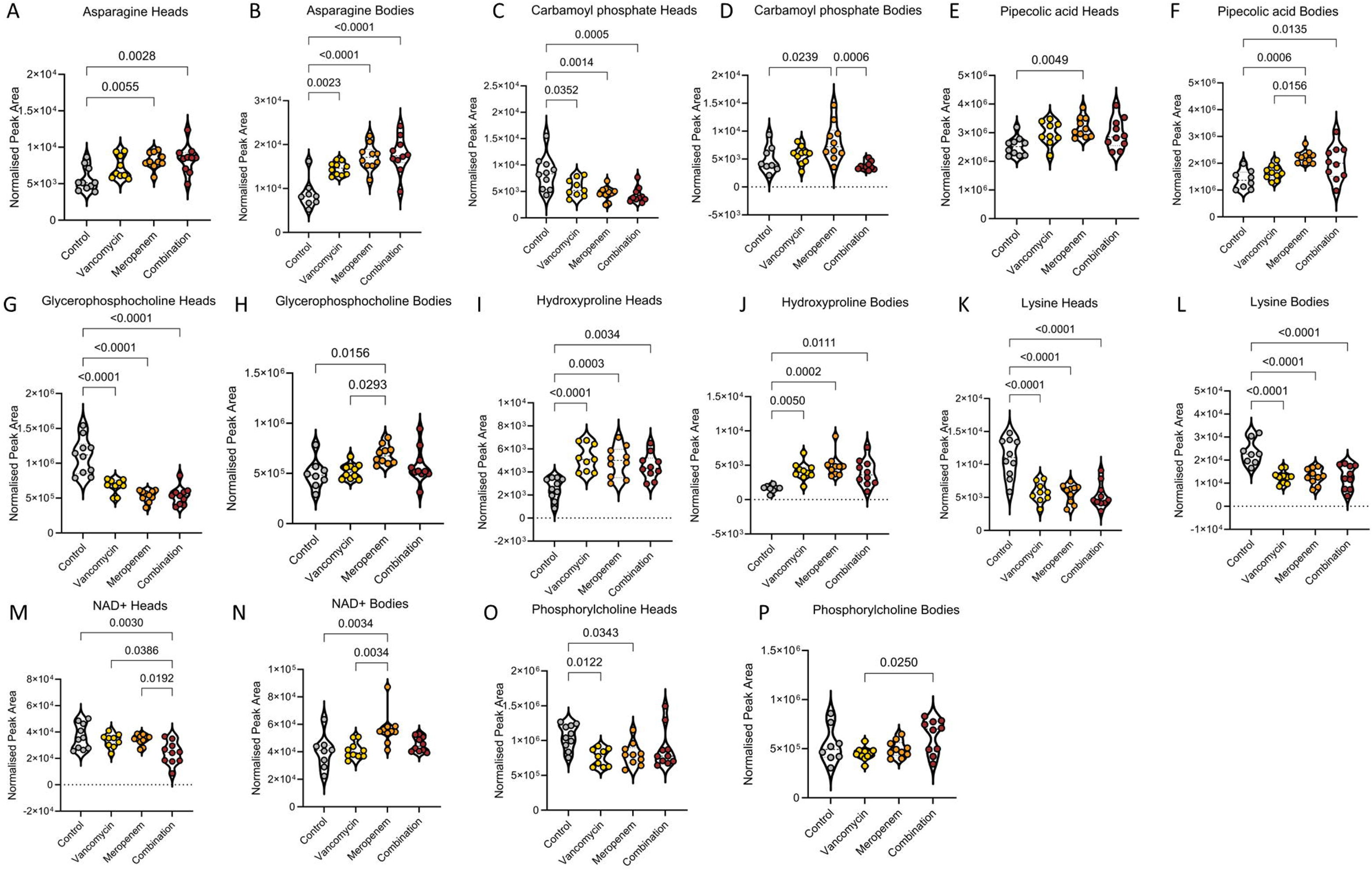
Antibiotic-induced metabolite alterations affecting both head and body compartments. Violin plots showing normalised peak areas of metabolites significantly altered in both head and body extracts of zebrafish larvae measured by metabolomics following treatment with vancomycin, meropenem, or vancomycin + meropenem (combination) compared to untreated larvae (control). (**A**) head asparagine, **(B)** body asparagine, (**C**) head carbamoyl phosphate, **(D)** body carbamoyl phosphate, (**E**) head pipecolic acid, **(F)** body pipecolic acid, (**G**) head glycerophosphocholine, **(H)** body glycerophosphocholine, (**I**) head hydroxyproline, **(J)** body hydroxyproline, (**K**) head lysine, **(L)** body lysine, (**M**) head NAD⁺, **(N)** body NAD⁺, and (**O**) head phosphorylcholine levels and **(P)** body phosphorylcholine levels. Each dot represents a pool of five larvae; violins indicate distribution density. One-way ANOVA followed by Tukey’s post hoc tests. n = 8-10 replicates per treatment group.

### Body-specific metabolic alterations in response to antibiotic treatment in zebrafish larvae

The metabolic profiling of body samples revealed that 27 of the detected neurotransmitters and metabolites were significantly altered in the periphery, but not in the brain. Among neurotransmitters we found that meropenem, but not vancomycin or combination treatment, increased peripheral (i.e. body) levels of GABA, glutamate and L-DOPA compared to control larvae (Fig. 6A-C), suggesting that effects on peripheral neurotransmitter pools were specific to meropenem. Other peripheral neurotransmitter levels were not affected by antibiotic treatment (Table S4). Apart from meropenem-induced peripheral neurotransmitter alterations, we found that meropenem also had the strongest effects on the other detected peripheral metabolites (Fig. 6, Fig. S2). Meropenem-induced metabolite changes included increased peripheral levels of the amino acids proline, valine, histidine, threonine, leucine/isoleucine and aspartate (Fig. 6D-G; Fig. S2). In addition, meropenem treatment significantly increased peripheral guanine levels compared to vancomycin treatment (Fig. 6H) and elevated cytosine concentrations compared to control (Fig. 6I). Metabolites involved in one-carbon metabolism and methylation pathways, such as S-adenosyl-L-methionine and S-adenosyl-L-homocysteine (SAH), were also significantly increased in meropenem-treated larvae (Fig. 6J,K), as was betaine, a methyl donor upstream of this pathway (Fig. S2), indicating disruptions in methylation capacity. Moreover, glutathione levels were significantly higher in response to meropenem (Fig. 6L), suggesting altered oxidative stress handling. Of these meropenem-associated changes, all except leucine/isoleucine, cytosine and SAH remained significant after correction. The combination treatment group also had strong effects on the peripheral metabolome. Compared to control, combination treatment was the sole intervention to augment peripheral alanine levels (Fig. 6M) and it also increased peripheral N-acetyl-glutamine levels (Fig. 6N), as did meropenem treatment. Moreover, combination treatment led to a rise of N-acetyl-alanine compared to all other groups (Fig. 6O) and a rise in proline and cytosine, as noted above (Fig. 6D, I). Additionally, inosine monophosphate (IMP), sarcosine and UDP-D-glucose, a metabolite linked to carbohydrate metabolism, were significantly increased in the combination only (Fig. 6P, Fig. S2). However, only proline, N-acetyl-glutamine and N-acetyl-alanine remained significant after correction.

**Figure 6:**
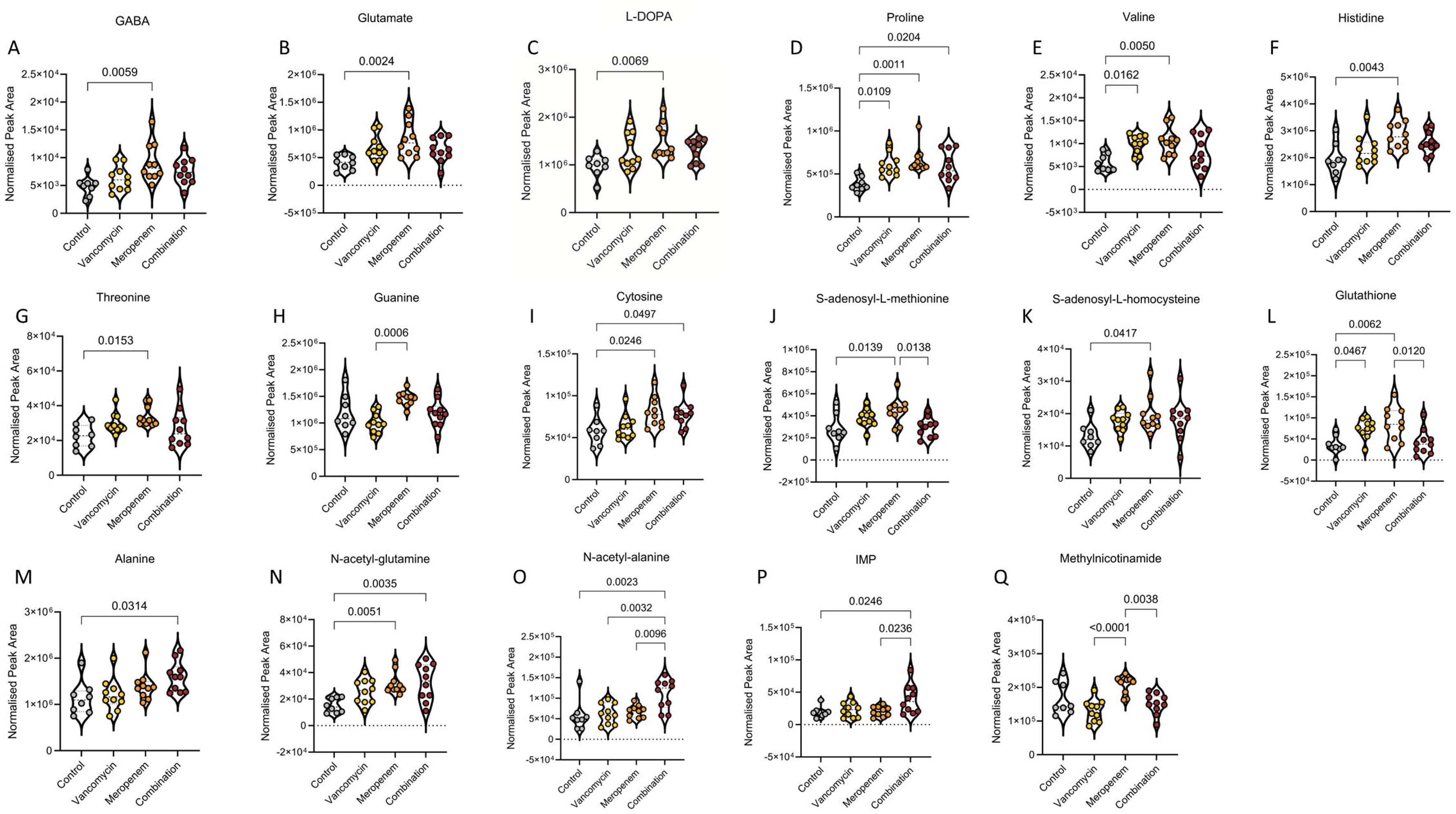
Body-specific metabolite alterations after larval antibiotic treatment. Violin plots showing normalised peak areas of neurotransmitters and metabolites measured by targeted and extended metabolomics, respectively, in body extracts of zebrafish larvae following treatment with vancomycin, meropenem, or vancomycin + meropenem (combination) compared to untreated larvae (control). (**A**) GABA, (**B**) glutamate, (**C**) L-DOPA, (**D**) proline, (**E**) valine, (**F**) histidine, (**G**) threonine, (**H**) guanine, (**I**) cytosine, (**J**) S-adenosyl-L-methionine, (**K**) S-adenosyl-homocysteine, (**L**) glutathione, (**M**) alanine, (**N**) N-acetyl-glutamine, (**O**) N-acetyl-alanine, (**P**) IMP, and (**Q**) methylnicotinamide. Each dot represents a pool of five larvae; violins indicate distribution density. One-way ANOVA followed by Tukey’s post hoc tests. n = 8-10 replicates per treatment group.

Notably, some metabolites such as IMP, S-adenosyl-L-methionine, glutathione, riboflavin and N-acetyl-alanine were significantly altered between the combination and the meropenem treatment group, suggesting that the addition of vancomycin modifies some of the metabolic effects induced by meropenem alone (Fig. 6J,L,O,P; Fig. S2). Vancomycin treatment itself induced fewer metabolic alterations that were confined to the periphery than meropenem and combination treatment. It led to a rise of peripheral proline, valine, aspartate and glutathione compared to controls, of which the rise in proline and valine remained significant after correction (Fig. 6D,E,L; Fig. S2). Differences between vancomycin and meropenem treatment were also evident in some peripheral metabolites such as pipecolic acid, methylnicotinamide, NAD^+^, glycerophosphocholine, p-hydroxybenzoate and guanine (Fig. 5F,H,N; Fig. 6H,Q; Fig. S2), of which all but glycerophosphocholine and p-hydroxybenzoate remained significant after correction, highlighting distinct peripheral metabolic impacts of the two antibiotics.

## Discussion

The microbiota-gut-brain axis is increasingly recognised as a key pathway through which microbial communities influence host neurodevelopment, metabolism, and behaviour (Cryan & Dinan, 2012; Sharon et al., 2016; Morais et al., 2021; Petracco et al., 2025). Using antibiotic-induced perturbation of the larval zebrafish microbiome, our study provides evidence that disruption of early-life microbial community structure is associated with behavioural alterations, changes in serotonergic signalling and widespread metabolic remodelling in the developing brain and gut. The three antibiotic treatments used in this study produced distinct microbial, behavioural and metabolic phenotypes, the magnitude of which did not simply scale with the extent of community disruption, indicating that the nature rather than the magnitude of microbiome alterations shapes host outcomes.

Early-life zebrafish microbiomes are typically dominated by Gram-negative *Proteobacteria* derived from the surrounding aquatic environment (Roeselers et al., 2011; Stephens et al., 2016; Garibay-Valdez et al., 2024), a pattern also observed in our data. Consistent with this, vancomycin had only limited effects on microbial community structure, because its main target Gram-positive bacteria show only very low abundance in the larval microbiome. Interestingly however, vancomycin treatment led to a shift within *Proteobacteria* depleting *Aeromonas* and increasing the relative abundance of *Pseudomonas*. Given that vancomycin cannot act on either genus directly (Levine, 2006), this likely reflects an indirect effect, for example competitive displacement of *Aeromonas* by *Pseudomonas* or altered host-microbe interactions following antibiotic exposure. In contrast, meropenem is known for its strong effects against Gram-negative bacteria including *Aeromonas*, *Pseudomonas* and *Achromobacter* (Zhanel et al., 2007), and therefore induced broader taxonomic restructuring. The depletion of the dominant taxa was accompanied by increased relative abundance of *Bacteroidota*, *Firmicutes* and *Actinobacteriota*, as well as of *Stenotrophomonas,* a Gammaproteobacterium whose intrinsic metallo-β-lactamase confers resistance to meropenem (Gil-Gil et al., 2020). Notably, alpha diversity increased after meropenem treatment, although antibiotic exposure is more often associated with reduced diversity. Such increases can occur when dominant taxa are removed, raising community evenness and allowing previously rare environmental microbes to be detected or to colonise (Dethlefsen & Relman, 2011; Zaneveld et al., 2017), a dynamic that is particularly plausible given the environmentally shaped nature of the early larval microbiota (Burns et al., 2016). The combined meropenem-vancomycin treatment produced a community restructuring similar to meropenem alone, with depletion of the dominant *Proteobacteria* taxa and enrichment of lineages with intrinsic β-lactam resistance mechanisms. This suggests that the addition of vancomycin had little additional effect on larval microbial community composition. An exception is the unique enrichment of the *Legionella* lineage in the combination group, possibly reflecting competitive release within the restructured community, as the combination group also showed a reduced relative expansion of *Stenotrophomonas*.

Consistent with a role of the microbiota in regulating host behaviour in our model system, antibiotic-treated larvae exhibited altered anxiety-like behaviour in the light-dark transition assay. Most notably, control larvae displayed robust increases in locomotion in the centre zone of the test arena during dark phases compared to light phases, an effect absent in larvae treated with meropenem/vancomycin combination and vancomycin alone. In contrast, meropenem-treated larvae showed a response similar to that of control animals, suggesting treatment-specific effects on this measure. When directly comparing dark phase locomotion in the centre zone, all antibiotic treatments reduced locomotion indicating an anxiogenic effect common to all groups. These treatment-specific behavioural effects cannot be directly explained by the microbiota alterations described above, because meropenem and combination treatment exerted both strong and similar effects on the larval microbiome, while vancomycin had only limited effects. Behaviourally, however, it was vancomycin and the combination that grouped together. Therefore, the behavioural effects might be a consequence of treatment-specific alterations in microbiota-host communication and/or distinct metabolic effects induced by the different treatments. Anxiety-related responses have been reported previously following a variety of microbiome manipulations in both zebrafish and mammalian models (Cryan et al., 2019; Kiran et al., 2024). In zebrafish larvae, microbiota disruption has been shown to influence behaviour and neurodevelopment as well. Specifically, 6 dpf germ-free larvae were found to be less anxious in a spontaneous free-swimming assay (Davis, Bryda, et al., 2016) and 10 dpf axenic and antibiotic-treated larvae (amphotericin B, kanamycin and ampicillin) were found to be hyperactive (Phelps et al., 2017). Together with our study, this suggests that various microbiota perturbations are able to alter larval behaviour in different ways necessitating a better understanding of the underlying mechanisms of each intervention.

When analysing the metabolite changes in response to the antibiotics used in this study, we found that many metabolite alterations were treatment-specific. Notably, meropenem monotreatment was the only intervention affecting neurotransmitter levels including highest central serotonin levels of all groups. Serotonin is known to regulate anxiety across vertebrates including zebrafish (Herculano & Maximino, 2014; Bacqué-Cazenave et al., 2020), suggesting one potential pathway by which meropenem affected behaviour in this group. Interestingly, several taxa enriched after meropenem treatment, including members of the *Bacteroidota* and *Firmicutes*, are capable of producing short-chain fatty acids and tryptophan derivatives that can influence host serotonin production, including in the brain (Clarke et al., 2013; Yano et al., 2015; Strandwitz, 2018). However, neither tryptophan nor indole, a bacterial tryptophan derivative, was altered by meropenem, and short-chain fatty acids were not measured, so such a link remains speculative. The gene expression data showed a similar treatment specificity, with the serotonin transporter *slc6a4a* reduced only after meropenem, while *tph1a*, encoding the rate-limiting enzyme of serotonin synthesis, was reduced by all three treatments. A reduction in *tph1a* would not necessarily be expected to alter total serotonin levels, as this isoform is expressed in the pineal organ and diencephalic subpopulations rather than in the raphe neurons supplying most of the brain (Norton et al., 2008; Lillesaar, 2011)

Beyond serotonergic signalling, meropenem treatment also increased peripheral glutamate, GABA, and L-DOPA concentrations. These changes may reflect altered microbial production or usage of neuroactive compounds in line with the PICRUSt2 prediction of reduced glutamine synthetase activity and amino acid transport capacity in the meropenem group. Specifically, reduced glutamine synthetase activity would spare glutamate from conversion to glutamine and thereby increase the substrate available for GABA synthesis (Otaru et al., 2021). It should be noted though that these predictions are inferred from taxonomic composition and are therefore not independent of the community changes described above. In addition, the altered neurotransmitter levels might also be the consequence of altered substrate availability or compensatory host responses to microbiome shifts, because numerous gut bacteria are capable of synthesising or modulating neurotransmitter precursors and metabolites (Strandwitz, 2018; Morais et al., 2021).

Vancomycin treatment did not alter neurotransmitter levels in larvae, but it was the only treatment which decreased arginine and N-acetyl-L-ornithine in larval heads, important contributors to the upstream stages of polyamine metabolism (Lillie et al., 2024). N-acetyl-L-ornithine is of particular interest here, as acetylated ornithine intermediates are characteristic of microbial arginine biosynthesis and are not generated by the vertebrate pathway (Xu et al., 2007), consistent with a microbial origin of this metabolite. Spermine, the terminal product of the pathway, was by contrast reduced by all three treatments. Polyamines are essential mediators of neuronal development, synaptic plasticity, and cellular stress responses (Minois et al., 2011; Hofer et al., 2021), and their depletion during early life may therefore carry consequences that extend beyond immediate metabolic disruption. Interestingly, two of the significant PICRUSt2 predictions were also related to polyamine transport capacity, but we did not detect differences in the peripheral levels of polyamine pathway metabolites. Vancomycin treatment was also the only treatment to reduce central biotin levels, a B vitamin produced by gut microbiota important for neural function (Karachaliou & Livaniou, 2024). Given that larval biotin derives from microbial synthesis and maternal yolk reserves (Quinlivan & Farber, 2017), this reduction may reflect impaired microbial production, reduced mobilisation from the yolk sac, or both. The finding is consistent with mammalian studies demonstrating that antibiotic-induced dysbiosis reduces host biotin levels through depletion of microbial biotin-producing taxa (Hayashi et al., 2017; Yoshii et al., 2019; Belda et al., 2022).

A notable unique metabolic effect of the combination treatment was a reduction in central NAD^+^ levels compared to all other groups, suggesting impaired redox balance and mitochondrial metabolic capacity within the developing brain (Covarrubias et al., 2021). In the periphery, the combination group also showed lower levels of S-adenosyl-L-methionine, glutathione and methylnicotinamide than meropenem-treated larvae, suggesting that the addition of vancomycin prevented the increase of methylation- and redox-related metabolites observed after meropenem alone. Since methylnicotinamide derives from nictotinamide, the precursor of NAD^+^ salvage, these effects point to altered NAD^+^ metabolism after combination treatment (Roberti et al., 2021). Together these observations indicate that dual antibiotic exposure modifies rather than simply amplifies microbiome-dependent metabolic perturbations.

Other metabolites were changed in the same manner by all antibiotic treatments in both head and body compartments including lysine, hydroxyproline and asparagine, pointing to broad disruption of amino acid and nitrogen metabolism independent of the antibiotic class used. The strong reduction of lysine after antibiotic treatment is particularly striking given that lysine is an essential amino acid not synthesised by vertebrates. In contrast, bacteria are able to synthesize lysine and in mammalian systems measurable microbial contributions to host lysine pools have been demonstrated (Metges et al., 1999; Neis et al., 2015; McCann & Rawls, 2023). However, in zebrafish larvae lysine is also derived from the yolk sac (Quinlivan & Farber, 2017), suggesting that the reduction may reflect impaired microbially-derived lysine uptake and/or reduced mobilization of yolk-derived lysine. Interestingly, asparagine and hydroxyproline concentrations increased in head and body samples in all treatment groups, alongside the reduction in lysine. These two observations may be connected, as cells respond to a shortage of essential amino acids by increasing asparagine synthesis from aspartate and glutamine (Balasubramanian et al., 2013), so that the loss of lysine could itself explain the accumulation of asparagine. The increased hydroxyproline levels might instead reflect altered collagen turnover and extracellular matrix remodelling, given that proline and hydroxyproline are the most abundant amino acids in collagen (Wu et al., 2011). Notably, the altered levels of lysine, hydroxyproline and asparagine may also reflect microbiome-independent effects of vancomycin and meropenem, as it cannot be excluded that the antibiotics acted directly on host tissues, although their chemical profile precludes absorption in the gastrointestinal tract (Levine, 2006; Zhanel et al., 2007)and uptake across skin and gills under immersion (Guarin et al., 2021). However, not all metabolites followed a consistent pattern across compartments. For example, carbamoyl phosphate and glycerophosphocholine were reduced in heads in all treatment groups but increased in bodies after meropenem. Such compartment-specific patterns argue against a uniform systemic action of the antibiotics on host tissues. Future studies using germ-free zebrafish, microbial recolonization, or microbiota transplantation approaches will therefore be important to disentangle microbiota-mediated effects from potential host-directed actions of antibiotics.

## Conclusion

In conclusion, this study demonstrates that antibiotic-induced disruption of the gut microbiome during early zebrafish larval development has strong consequences for host behaviour, neurotransmitter signalling, and metabolism. Using vancomycin, meropenem and their combination as tools to restructure the microbial community, we show that the nature of the disruption, rather than its extent, shaped neurochemical and metabolic outcomes. Behavioural analyses revealed both shared and treatment-specific alterations of anxiety-like behaviour accompanied by reduced *tph1a* expression in all treatment groups, changes in peripheral neurotransmitters after meropenem and in central polyamine metabolism after vancomycin. Extended metabolomics further revealed that larval antibiotic treatment disrupts a broad network of metabolic pathways in both head and body compartments, including amino acid homeostasis, redox and methylation balance and microbiome-derived cofactor supply. Establishing which microbial taxa and metabolic outputs drive these host phenotypes will require future targeted microbial reconstitution experiments. Nonetheless, our results underscore the importance of considering the microbial, neurochemical and metabolic consequences of early-life antibiotic exposure, with potential implications for understanding how microbiome perturbation during development may contribute to lasting changes in brain function and behaviour.

## Methods

### Experimental animals and housing conditions

Experiments were performed with 5 dpf zebrafish larvae (*Danio rerio*, TU strain) obtained from the local breeding colony maintained at the Medical University of Graz. For embryo production, adult TU wild-type zebrafish were kept overnight in breeding tanks at a 1:1 sex ratio (three males and three females per tank) at 28°C under a 14/10h light/dark cycle. After spawning on the following day, the eggs were collected and rinsed in autoclaved 1x E3 medium (5mM NaCl, 0.17mM KCl, 0.33mM CaCl₂ ⋅ 2H₂O, 0.33mM MgSO₄). The fertilized eggs were incubated in Petri dishes in autoclaved 1x E3 medium at 28°C medium temperature in an incubator (Memmert, Schwabach, Germany) with a 14/10h light/dark cycle. At 3 days post fertilization (dpf) the hatched and morphologically normal larvae were selected for experiments. An equal number of larvae from 2-4 different breeding tanks were pooled and transferred into fresh petri dishes filled with 60mL autoclaved 1x E3 medium (30 larvae per dish) for subsequent antibiotic treatment. The zebrafish facility features optimal housing conditions for zebrafish (Tecniplast, Buguggiate, Italy), a 14-hour light/10-hour dark cycle, standard feeding schedules (Gemma fish feed twice a day and artemia once a day) and dedicated personnel. Animal experiments were approved by the Austrian Federal Ministry of Women, Science and Research (BMFWF) under the project licence (GZ: 2026-0.032.315).

### Antibiotic exposure

The antibiotic concentrations were selected based on previous studies in zebrafish embryos, larvae or adults (Geiger et al., 2013; Dinday & Baraban, 2015; Kato et al., 2020; Habjan et al., 2021; Lin et al., 2023). At 3 dpf zebrafish larvae were exposed to 100 µM vancomycin (Thermo Fisher Scientific), 1 mM meropenem (Fresenius Kabi) or a combination of both at the same concentrations as in the single-treatment groups. The antibiotics were dissolved in autoclaved E3 medium and administered to the larvae in the treatment groups. Control animals were kept in autoclaved E3 medium without antibiotics. The medium was replaced after 24 hours treatment at 4 dpf in all groups to keep the antibiotic concentrations constant during the exposure period. The larvae were kept in an incubator at 28°C with a 14/10h light/dark cycle for a total exposure period of 48h until 5 dpf. No dead or abnormal larvae were observed during the treatments.

### Behavioural testing

#### Light-Dark (LD) Transition Assay

At 5 dpf, one cohort of larvae from antibiotic-treated and control groups were subjected to a standardized LD transition assay in a Zantiks MWP unit (Zantiks Ltd, UK) adapted from (Hillman et al. 2024). Larvae (n = 15 per group) were individually transferred into sterile 48-well plates using sterile Pasteur pipettes, ensuring minimum handling stress. The residual medium was carefully removed and replaced with 1 mL of fresh autoclaved E3 medium. Plates with a well diameter of 10.38 mm were loaded into the Zantiks behavioural system and acclimatized for 30 min under constant white light at 28 °C (habituation phase). The assay consisted of three alternating 5-min cycles of light (350 lux) and dark (0 lux), for a total of 30 min (light-dark-light-dark-light-dark) directly after the habituation phase. Locomotor activity in the inner (6.908 mm diameter) and outer zone of the test arena were automatically tracked using the integrated Zantiks software. The dark flash response was evaluated by quantifying distance moved in 1 second bins over the 10 seconds preceding and the 10 seconds following the first light/dark transition. Following testing, larvae were euthanized and collected for 16S rRNA sequencing.

### 16S rDNA amplicon sequencing and microbial community analysis

Microbial DNA was purified from 5 dpf whole zebrafish larvae by using a spin-column-based kid following manufacturers’ instructions (PureLink™ Microbiome DNA Purification Kit, Invitrogen). Six whole larvae were pooled per sample to obtain sufficient DNA for downstream 16S rRNA gene sequencing. Amplicon sequencing of the V3–V4 region of the bacterial 16S rRNA gene was performed by Novogene using an Illumina paired-end sequencing platform. Sequencing generated between approximately 34,000 and 168,000 raw paired-end reads per sample. Following read merging, quality filtering, and chimera removal, the resulting effective tags had an average length of approximately 410–427 bp. Paired-end sequencing was conducted, generating 250 bp reads from each end (FW: 5’-TCGTCGGCAGCGTCAGATGTGTATAAGAGACAGCCTACGGGNGGCWGCAG-3’, REV: 5’-GTCTCGTGGGCTCGGAGATGTGTATAAGAGACAGGACTACHVGGGTATCTAATCC-3’). Sequencing data were processed using the QIIME2 workflow (version 2022.02). Briefly, raw sequencing reads were quality-filtered, and denoising, paired-end merging and chimera removal were performed using DADA2 to generate amplicon sequence variants (ASVs) (Callahan et al., 2017). Taxonomic assignment of representative ASV sequences was conducted using the SILVA database (version 138.1). ASV tables were used to calculate relative abundance at different taxonomic levels. Alpha diversity indices (e.g., Shannon, Simpson, Chao1) were calculated to evaluate species richness and evenness. For beta diversity analysis, multiple sequence alignment of representative ASV sequences was performed and phylogenetic trees were constructed. Community structure differences among groups were assessed using principal coordinate analysis (PCoA). Differential abundance analysis between groups was performed using LEfSe to identify significantly enriched or depleted taxa. Metagenomic functional content was predicted from the ASV table using PICRUSt2 (R package V2.3.0) and annotated against the Kyoto Encyclopedia of Genes and Genomes (KEGG) Orthologues (KOs). Predicted KO and pathway were compared between groups using the Wilcoxon rank-sum test and corrected with Benjamini-Hochberg. Functional annotation results were integrated with taxonomic profiles to infer potential differences in microbial metabolic capacity among experimental groups.

### Real-time qPCR

A second cohort of zebrafish larvae was used for RNA analyses. Total RNA from 5 dpf zebrafish larvae heads was extracted using TRIzol™ reagent (Thermo Fisher Scientific) according to the manufacturer’s protocol followed by DNase treatment with the DNA-free™ DNA Removal Kit (Thermo Fisher Scientific). Each biological replicate consisted of a pool of five larval heads. 0.3 µg RNA was reverse-transcribed to cDNA using the High Capacity cDNA Reverse Transcription Kit (Thermo Fisher Scientific). The expression of *mao*, *slc6a4a*, *slc6a4b*, *tph1a* and *tph2* was analysed (Table S5). To evaluate the gene expression at the transcriptional level a real-time qPCR was performed using SsoAdvanced Universal SYBR Green Supermix (BIO-RAD) on a CFX Connect Real-Time PCR System (BIO-RAD), using the SYBR green detection protocol. Primers for genes of interest were taken from a previous publication (Reichmann et al., 2020). Amplification of the correct amplicon was confirmed by Sanger sequencing. All samples were measured in technical triplicate. *actin beta1* (*actb1*) and *ribosomal protein L13a* (*rpl13a*) were used as reference genes for quantification of target gene expression. Quantitative measurements of target gene levels relative to controls were performed with the ΔΔCq method using the mean ΔCq of the control group as the calibrator (Reichmann et al., 2020). Group differences were expressed as fold changes.

### Metabolomics and neurotransmitter quantification

For metabolomics analysis, larvae from a third cohort of animals were separated into heads and bodies by decapitation posterior to the operculum and 5 heads and 5 bodies were pooled per sample and frozen on dry ice. Frozen tissue samples were processed following Chetwynd et al., 2017 with minor modifications. Briefly, samples were homogenized in 50% MeOH-H₂O using Power Bead tubes and a PowerLyzer homogenizer. Polar metabolites were analysed using liquid chromatography couple to tandem mass spectrometry following the methodology described by (Yuan et al., 2012). We used an XBridge Amide column (100 mm x 2.1 mm x 4.6 µm, Waters, Manchester, UK), and separation was achieved with a Nexera UHPLC (SHIMADZU, Kyoto, Japan) with a gradient elution. Solvent additives for HILIC high-pH were ammonium acetate and 0.1% ammonium hydroxide. We used a SCIEX QTRAP 6500 triple quadrupole mass spectrometer (Applied Biosystems, Framingham, MA, USA) for detecting metabolites in both positive and negative polarity mode. Raw files were imported into Skyline software for peak integration. The resulting data was further normalized and statistically analysed using the online platform MetaboAnalyst. Protein concentrations of individual samples were determined using the Bradford protein assay (Thermo Scientific, Pierce^TM^ Bradford Plus Protein Assay Kit) for downstream analyses.

### Statistics

Statistical analysis was performed with GraphPad Prism 10.4.1 (Systat Software Inc., San Jose, USA), unless stated otherwise. Data are presented as mean ± SEM unless otherwise stated. A p-value < 0.05 was considered statistically significant. Behavioural data were analysed by repeated-measures ANOVA with Greenhouse-Geisser correction for non-sphericity, followed by Tukey’s post hoc multiple comparisons test for the readouts per-minute locomotion across the assay and second-by-second locomotion around the first light-to-dark transition. Average distance moved per phase and average inner-zone distance moved during light and dark phases were analysed by two-way ANOVA with light condition and treatment as factors, followed by Tukey’s post hoc multiple comparisons test. Gene expression data, neurotransmitter data, alpha diversity and metabolomics data were analysed by one-way analysis of variance (ANOVA) followed by Tukey’s post hoc multiple comparisons test. For neurotransmitter measurements, normalised peak areas were log2-transformed, and treatment-group differences were assessed separately for each metabolite using one-way ANOVA followed by Tukey’s multiple-comparisons test. For the extended metabolomics data, a secondary analysis was performed to assess robustness to multiple testing. Specifically, normalised peak areas were log2-transformed, all pairwise contrasts between treatment groups were estimated per metabolite using estimated marginal means, and Benjamini-Hochberg false discovery rate correction was applied globally across all comparisons within each compartment (85 metabolites × 6 pairwise comparisons). Comparisons remaining significant at q < 0.05 are indicated in Tables S2 and S3. Beta diversity was calculated using Bray-Curtis dissimilarity as well as weighted and unweighted UniFrac distances and analysed by permutational multivariate analysis of variance (PERMANOVA). Differentially abundant taxa were identified using linear discriminant analysis effect size (LEfSe), which applies a Kruskal-Wallis test followed by Wilcoxon rank-sum testing and linear discriminant analysis to estimate effect size (LDA score threshold: 4). In addition, pairwise LEfSe comparisons were performed between individual treatment groups.

## Acknowledgments

We are grateful to Sabine Donner and the team of the biomedical research facility of Medical University of Graz for technical assistance and animal care. We thank the European Zebrafish Resource Center (EZRC) of the Karlsruhe Institute of Technology (KIT) for providing the TU zebrafish line used in this study. This research was funded fully or in part by the Austrian Science Fund (FWF) [grant DOI: 10.55776/P35774 and grant DOI: 10.55776/PAT5589324] and the Medical University of Graz. For open access purposes, the author has applied a CC BY public copyright license to any author accepted manuscript version arising from this submission. AG is funded by the PhD program Molecular Medicine of the Medical University of Graz and IF is funded by the FWF within the PhD program Molecular Medicine of the Medical University of Graz.

