## Supplementary File for "Early-life vancomycin and meropenem treatment exert distinct effects on microbiota-gut-brain axis signalling in zebrafish larvae"

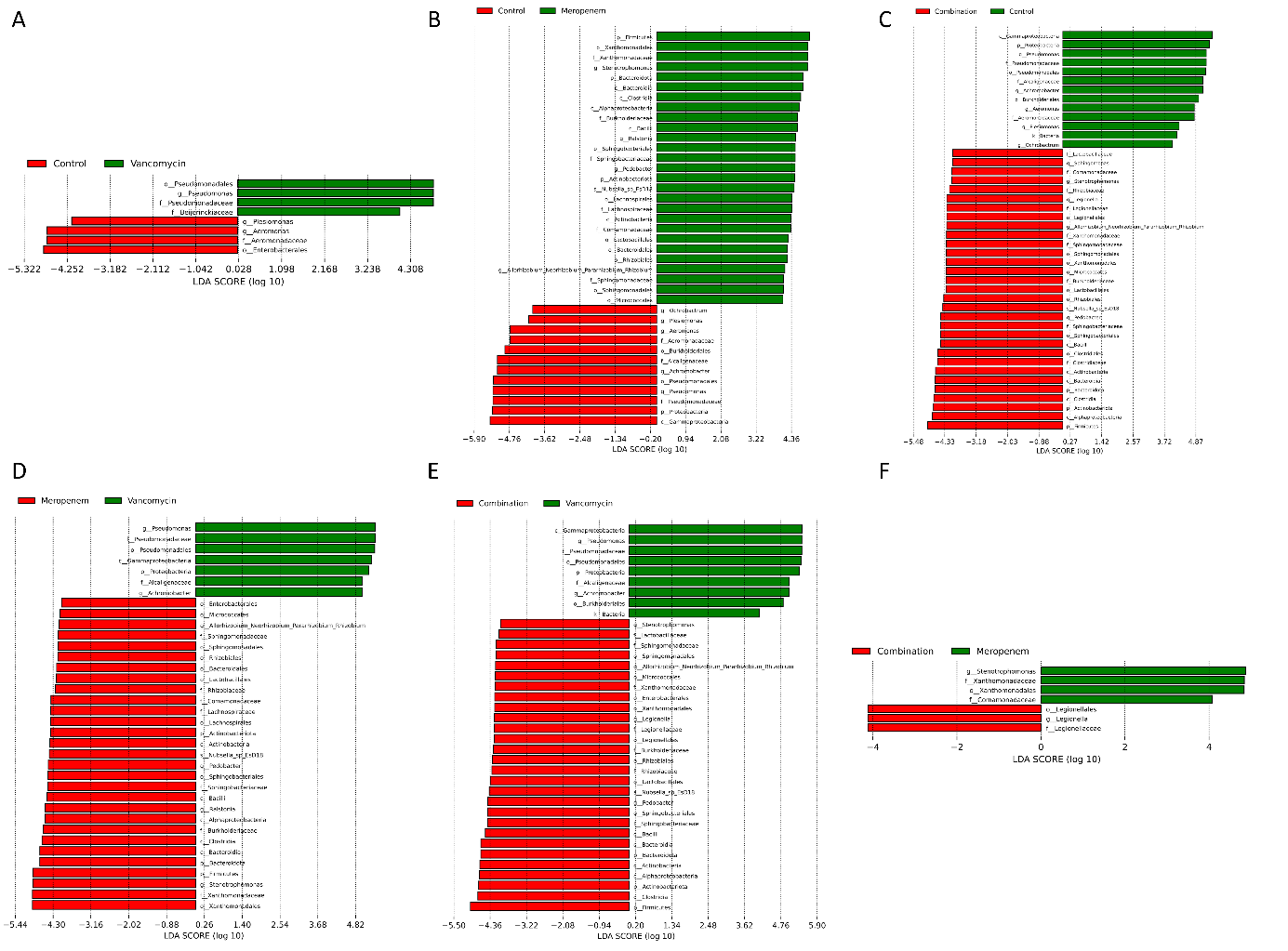

**Figure S1: Pairwise LEfSe analysis of differentially abundant taxa between treatment groups.** Linear discriminant analysis (LDA) scores for taxa differing significantly between (**A**) control and vancomycin, (**B**) control and meropenem, (**C**) control and combination, (**D**) vancomycin and meropenem, (**E**) vancomycin and combination, and (**F**) meropenem and combination treated larvae. Positive and negative LDA scores indicate enrichment in the respective group as indicated by the colour legend in each panel. Analysis was performed using LEfSe (LDA threshold 4). n = 10 replicates per group.

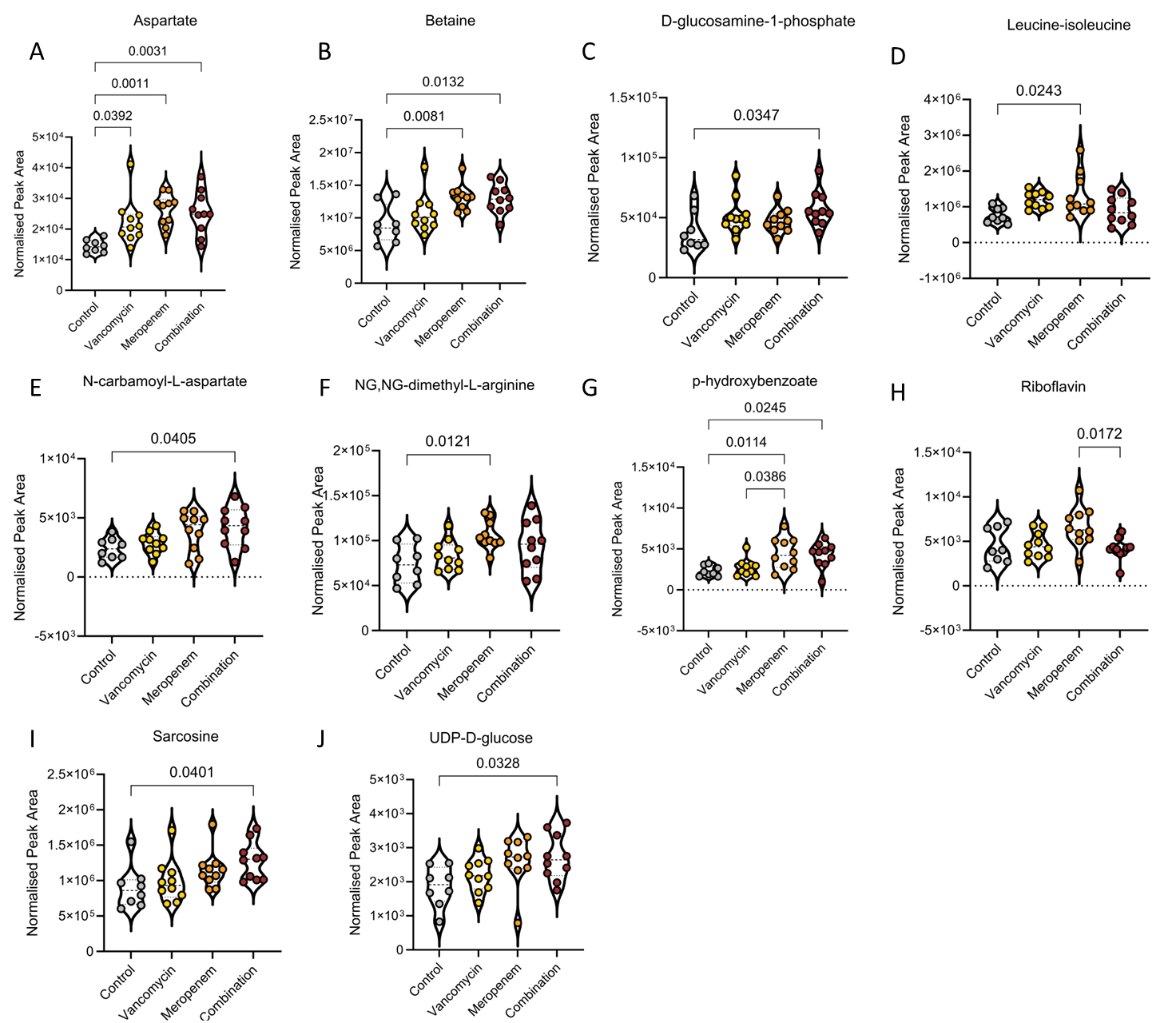
**Figure S2: Further metabolite alterations in body extracts of antibiotic-treated zebrafish larvae.** Violin plots showing normalised peak areas of metabolites significantly altered in body extracts of zebrafish larvae measured by extended metabolomics following treatment with vancomycin, meropenem, or vancomycin + meropenem (combination) compared to untreated larvae (control). (**A**) aspartate, (**B**) betaine, (**C**) D-glucosamine-1-phosphate, (**D**) leucine/isoleucine, (**E**) N-carbamoyl-L-aspartate, (**F**) NG,NG-dimethyl-L-arginine, (**G**) p-hydroxybenzoate, (**H**) riboflavin, (**I**) sarcosine and (**J**) UDP-D-glucose. Each dot represents a pool of five larvae; violins indicate distribution density. One-way ANOVA followed by Tukey's post hoc tests. n = 8–10 replicates per treatment group.

**Table S5: Primer sequences for qPCR.** Primer sequences for *monoamine oxidase* (*mao*), *solute carrier family 6 member 4a* (*slc6a4a*), *solute carrier family 6 member 4b* (*slc6a4b*), *tryptophan hydroxylase 1a* (*tph1a*), and *tryptophan hydroxylase 2 (tph2*).

| **Gene** | **Sequence forward primer (5’-3’)** | **Sequence reverse primer (5’-3’)** |
| --- | --- | --- |
| *mao* | CAG TTT GGC AGG GTC TTG CG | ATG GAG TTT GCC CAT TGC AC |
| *slc6a4a* | ACA ACC GAT GGA ACA CTC CC | CAA CAC CTG CCG GAC ATA AA |
| *slc6a4b* | GGA GAC CAG CGT ATG GGT AGA | CCA GAC TGG TCA CGA TAG CA |
| *tph1a* | TCT ACA CAC CTG AGC CAG AC | CCC TTC CTG CTT ACA GAG CC |
| *tph2* | TGG CTC TGA ACT GGA TGC TG | GGG CTG CCC AAA TTT GTA GT |
